# Naturally mutated spike proteins of SARS-CoV-2 variants show differential levels of cell entry

**DOI:** 10.1101/2020.06.15.151779

**Authors:** Seiya Ozono, Yanzhao Zhang, Hirotaka Ode, Toong Seng Tan, Kazuo Imai, Kazuyasu Miyoshi, Satoshi Kishigami, Takamasa Ueno, Yasumasa Iwatani, Tadaki Suzuki, Kenzo Tokunaga

**Author notes:** Correspondence: Kenzo Tokunaga, Department of Pathology, National Institute of Infectious Diseases, Shinjuku, Tokyo 162-8640, Japan. These authors contributed equally to this work (Seiya Ozono, Yanzhao Zhang, Hirotaka Ode).

## Abstract

The causative agent of the coronavirus disease 2019 (COVID-19) pandemic, severe acute respiratory syndrome coronavirus 2 (SARS-CoV-2), is steadily mutating during continuous transmission among humans. Such mutations can occur in the spike (S) protein that binds to the angiotensin-converting enzyme-2 (ACE2) receptor and is cleaved by transmembrane protease serine 2 (TMPRSS2). However, whether S mutations affect SARS-CoV-2 infectivity remains unknown. Here, we show that naturally occurring S mutations can reduce or enhance cell entry via ACE2 and TMPRSS2. A SARS-CoV-2 S-pseudotyped lentivirus exhibits substantially lower entry than SARS-CoV S. Among S variants, the D614G mutant shows the highest cell entry, as supported by structural observations. Nevertheless, the D614G mutant remains susceptible to neutralization by antisera against prototypic viruses. Taken together, these data indicate that the D614G mutation enhances viral infectivity while maintaining neutralization susceptibility.

---

The outbreak of coronavirus disease 2019 (COVID-19) caused by severe acute respiratory syndrome coronavirus 2 (SARS-CoV-2) (*1-4*) has rapidly spread around the globe from Wuhan, China, affecting more than 180 countries. During the COVID-19 pandemic, SARS-CoV-2 has accumulated mutations throughout viral genes encoding the ORF1a, ORF1b, ORF3, ORF8, nucleocapsid (N), and spike (S) proteins, etc., some of which are clade defining (Nextstrain, http://nextstrain.org/ncov, based on the GISAID data, www.gisaid.org). Mutations in the S protein are especially crucial because the S protein is key for the first step of viral transmission, i.e., entry into the cell by binding to the angiotensin-converting enzyme-2 (ACE2) receptor (*1*), followed by cleavage with transmembrane protease serine 2 (TMPRSS2) (*5,6*), both of which are abundantly expressed in not only the airways, lungs, and nasal/oral mucosa (*7,8*) but also the intestines (*9*).

To quantify S-mediated cell entry, we employed a novel measurement system for viral entry, which we have recently established by generating a lentiviral vector harboring a small luminescent peptide tag, HiBiT, allowing us to precisely normalize input virus doses. This system results in enhanced experimental accuracy in a single-round replication assay (*10*) in which a small but detectable difference in cell entry on a linear scale is critical because such a difference can be amplified in multiple virus replication cycles. By using this system, we first examined the viral entry activity of lentiviruses pseudotyped with SARS-CoV (Urbani strain) S or SARS-CoV-2 (Wuhan-Hu-1 strain) S (referred to hereafter as SARS-S and SARS2-S, respectively) in cells expressing ACE2, a major receptor for SARS-S (*11*). The lentivirus pseudotyped with SARS-S was able to efficiently enter 293T cells expressing ACE2, whereas the SARS2-S-pseudotyped virus entered the cells much less efficiently (Fig. S1). This result suggests that the expression of ACE2 alone is insufficient to support cell entry of SARS-CoV-2.

Because accumulated evidence has shown that the expression of TMPRSS2 enhances both SARS-CoV and SARS-CoV-2 infection (*5,6*), we next tested whether the coexpression of ACE2 and TMPRSS2 could mediate efficient infection of 293T cells with SARS2-S-pseudotyped lentiviruses. Notably, the dual expression of ACE2 and TMPRSS2 markedly facilitated SARS2-S-mediated cell entry while moderately enhancing entry by the SARS-S pseudovirus (Fig. 1A). These results indicate that SARS2-S-mediated entry into cells is more highly dependent on TMPRSS2 coexpression than that of SARS-CoV, suggesting that they may have different cellular tropisms to some extent.

**Figure 1.**
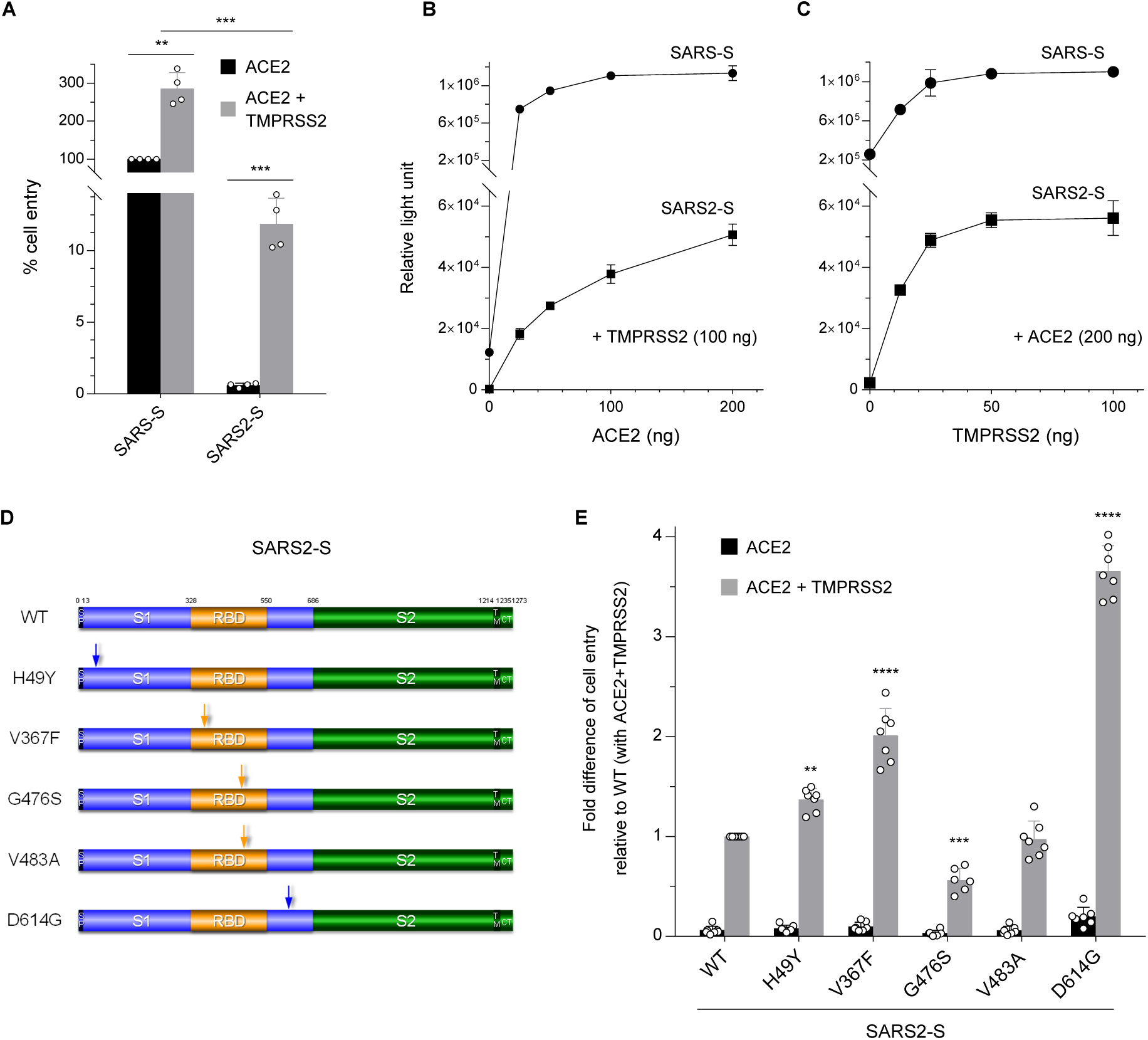
The SARS-CoV-2 S protein has significantly low cell entry activity, with a strong dependency on TMPRSS2, and its variant proteins display differential levels of entry activity. (A) Viruses were prepared by transfection of 293T cells with the HiBiT-tagged lentiviral packaging plasmid, the firefly luciferase-reporter lentiviral transfer plasmid, and either a SARS-CoV S (SARS-S) or SARS-CoV-2 S (SARS2-S) expression plasmid. Viral supernatants were subjected to HiBiT assays, and S-pseudotyped viruses normalized by HiBiT activity were used for infection of 293T cells expressing the host receptor ACE2 alone (black) or coexpressing TMPRSS2 (gray). Cell entry was determined by firefly luciferase activity in cell lysates. Data from four experiments are shown as a percentage of cell entry of SARS-S-pseudotyped viruses into 293T cells expressing ACE2 only (mean ± s.d., *n* = 3 technical replicates). The *p* value was calculated using two-tailed paired Student’s *t*-test, \*\**p* < 0.005, \*\*\**p* < 0.001. (B, C) The effect of ACE2 or TMPRSS2 expression levels on cell entry activity. 293T cells were transfected with a high and constant level of an expression plasmid encoding ACE2 together with increasing levels of a TMPRSS2 expression plasmid (*B*), and *vice versa* (*C*). Transfected cells were infected with lentiviruses pseudotyped with either SARS-S (circle) or SARS2-S (square), as described in *A*. Data shown are representative of three independent experiments (mean± s.d., *n* = 3 technical replicates). (D) Schematic illustration of the prototype (wild-type, WT) and globally spread variant SARS2-S proteins. Numbers indicate amino acid positions. The signal peptide (SP), transmembrane domain (TM), cytoplasmic tail (CT), S1 subunit (S1), S2 subunit (S2), and receptor-binding domain (RBD) are indicated. The positions of mutations are indicated by arrows. (E) Functional comparison of the entry activity of WT and mutant SARS2-S proteins. Different S-pseudotyped viruses were prepared as described in *A* and used for infection of 293T cells expressing the host receptor ACE2 alone (black) or coexpressing TMPRSS2 (gray). Cell entry was determined by firefly luciferase activity in cell lysates. Data are shown as the fold difference of cell entry relative to that of WT into 293T cells coexpressing ACE2 and TMPRSS2 (mean± s.d. from six to seven independent experiments with three technical replicates); \*\**p* < 0.005, \*\*\**p* < 0.001, \*\*\*\**p* < 0.0001 compared with WT using one-way ANOVA with Dunnett’s multiple comparison test.

Nevertheless, ∼25-fold differences in cell entry between SARS-S and SARS2-S pseudoviruses were observed (Fig. 1A), leading to the hypothesis that SARS2-S might be less incorporated into lentiviral particles than SARS-S, probably due to reduced compatibility with lentiviral particles. Compared with SARS-S, SARS2-S has one (alanine-to-cysteine at position 1247) and two (valine-to-isoleucine at position 1216 and leucine-to-methionine at position 1233) amino acid differences in the cytoplasmic tail (CT) and transmembrane (TM) domain, respectively (Fig. S2A). Thus, we created a SARS2-S C1247A mutant and a chimeric SARS2-S harboring the TM/CT domains of SARS-S. All SARS2-S proteins showed comparable levels of cell entry (Fig. S2B) and actual virion incorporation (Fig. S2C). These results indicate that the significantly lower rate of cell entry of the SARS2-S pseudovirus was not due to differences in the efficiency of S protein incorporation into virions but rather to the intrinsic nature of the SARS2-S protein.

To further assess differences between the SARS-S and SARS2-S proteins, we next addressed whether these S proteins might differ in their ability to utilize a given level of cell surface ACE2 or TMPRSS2. Based on lentiviral infection of 293T cells expressing a high and constant level of ACE2 together with a range of expression levels of TMPRSS2 and *vice versa*, SARS2-S-mediated infection required higher levels of cell-surface ACE2 and TMPRSS2 expression than SARS-S to attain maximum levels of infectivity (Fig. 1B and 1C). Therefore, it is likely that SARS2-S is less adapted to ACE2 due to the inefficient usage of this host protein and essentially requires sufficient levels of TMPRSS2 expression.

Next, we investigated using our assay system whether naturally occurring mutations in the S protein affect the cell entry of SARS-CoV-2 We created plasmids expressing five different S variants that were initially identified in China (H49Y (*12*)), Europe (V367F (*13*) and D614G (*14-16*)), and the United States (G476S and V483A (*13*)) (Fig. 1D) and examined the effects of these mutations on entry into cells expressing ACE2 and TMPRSS2, by comparison with that of wild-type (WT) S protein. These naturally occurring S mutations resulted in reduced (G476S), equal (V483A), or enhanced (D614G, V367F, and H49Y) cell entry. Remarkably, the D614G mutant displayed the highest level of entry activity compared with that of WT S protein (Fig. 1E). This result is particularly important because the D614G mutation defines the clade A2a (also called G) that is rapidly spreading worldwide, accounting for the great majority of isolates (*17-20*).

To analyze differences among WT/mutant SARS2-S and SARS-S, we performed structural analyses on complex models between ACE2 and these S proteins (Fig. 2A and 2B). Interestingly, the SARS-S trimer showed an open conformation, providing a larger contact area in the receptor-binding domain (RBD) that specifically interacts with ACE2; this might lead to higher accessibility of the SARS-S RBD to ACE2 than that of SARS2-S. This finding is indeed consistent with the recent report that the SARS-CoV-2 S protein has lower ACE2 binding affinity than SARS-CoV S (*21*). It is also likely that to a large extent, the structures of SARS2-S mutants reflect differential cell entry (Fig. 2C; see details in the legend). Notably, the aspartic acid residue at position 614 located in the S1 subunit of the WT protein (D614) is able to form a hydrogen bond with a threonine residue at position 859 (T859), as recently reported (*19*), and/or a salt bridge with a lysine residue at position 854 (K854) located in the S2 subunit of the other protomer (Fig. 2D). This finding suggests in turn that the mutation of this residue to a glycine (G614) can provide flexible space between two protomers due to the short side chain, allowing the S1 subunit to be dissociated more smoothly from the S2 subunit, and/or likely providing conformational flexibility to the overall structure of the S trimer, which might lead to improved accessibility of ACE2 into the RBD. These results are fully consistent with the high entry activity of the virus harboring this mutation (Fig. 1E).

**Figure 2.**
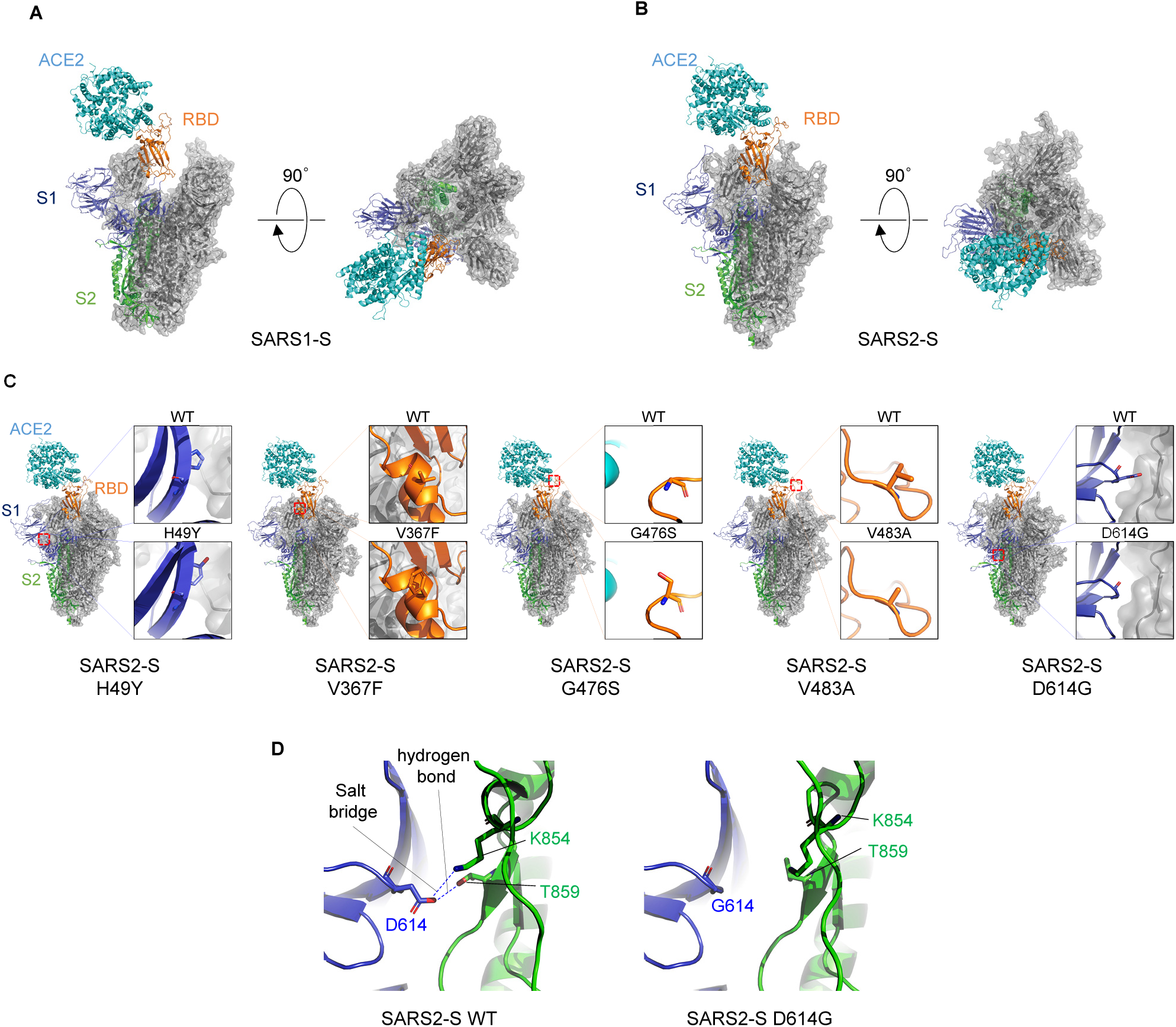
Structural comparison of S proteins. (A, B) Trimeric structure of the S proteins from SARS-CoV *(A*) and SARS-CoV-2 (*B*) that bind to the host receptor ACE2 (cyan). One protomer of the S protein is shown as a ribbon in navy (S1 subunit), orange (receptor-binding domain (RBD) in S1 subunit), and green (S2 subunit), while two other protomers are shown as gray transparent surfaces and ribbons. The structures are viewed from two different angles. (C) Comparison between wild-type (WT) and mutant S proteins from globally spread SARS-CoV-2 variants. (*Inset*) Enlargement of the region in which each amino acid is mutated, with comparison with the WT S protein. *H49Y*; as the histidine at position 49 is located distant from the RBD and putative cleavage sites, the effect of this mutation on S’s function is likely limited. *V367F*; the substitution from a valine to a phenylalanine at position 367 in the RBD introduces a larger side chain at a protomer–protomer interface, which might provide a more rigid RBD structure. *G476S*; the substitution at position 476 in the RBD results in a protruded surface, which appears to interfere with the ACE2-RDB interaction. *V483A*; both valine and alanine residues have short side chains, likely sharing similar phenotypes. *D614G*; details are depicted in *Fig. 2D*. Note that these structural bases are largely consistent with the results of cell entry activity shown in *Fig. 1D*. (D) Structural difference between WT and D614G SARS-CoV-2 S proteins. *WT* (left); an aspartic acid (D614) in the S1 subunit (navy) of a protomer binds to a threonine (T859) and/or a lysine (K854) in the S2 subunit (green) of the other protomer though electrostatic interaction between the pairs of these residues. *D614G* (right); the short nonpolar side chain of glycine (G614), which does not bind to T859 and K854, provides flexible space between the two protomers. The figures were drawn with PyMOL ver. 2.4 (https://pymol.org).

Given that the D614G S protein is biologically and structurally different from the WT protein, we hypothesized that this mutation might affect the antigenicity of the S protein. To examine this possibility, we performed neutralization assays to compare the neutralizing sensitivity of the WT and D614G S proteins to anti-SARS-CoV-2 sera. For the neutralization assays, we utilized serum samples derived from five patients confirmed to be infected with prototype viruses and a control serum from a healthy donor. Regardless of serum concentration, the anti-SARS-CoV-2 patient sera but not the control serum efficiently neutralized both viruses pseudotyped with the SARS2-S WT and D614G mutant proteins (Fig. 3). These results indicate that the D614G mutation in the SARS2-S protein maintains neutralization sensitivity to the anti-SARS2-S antibodies, *i*.*e*., its antigenicity *per se*.

**Figure 3.**
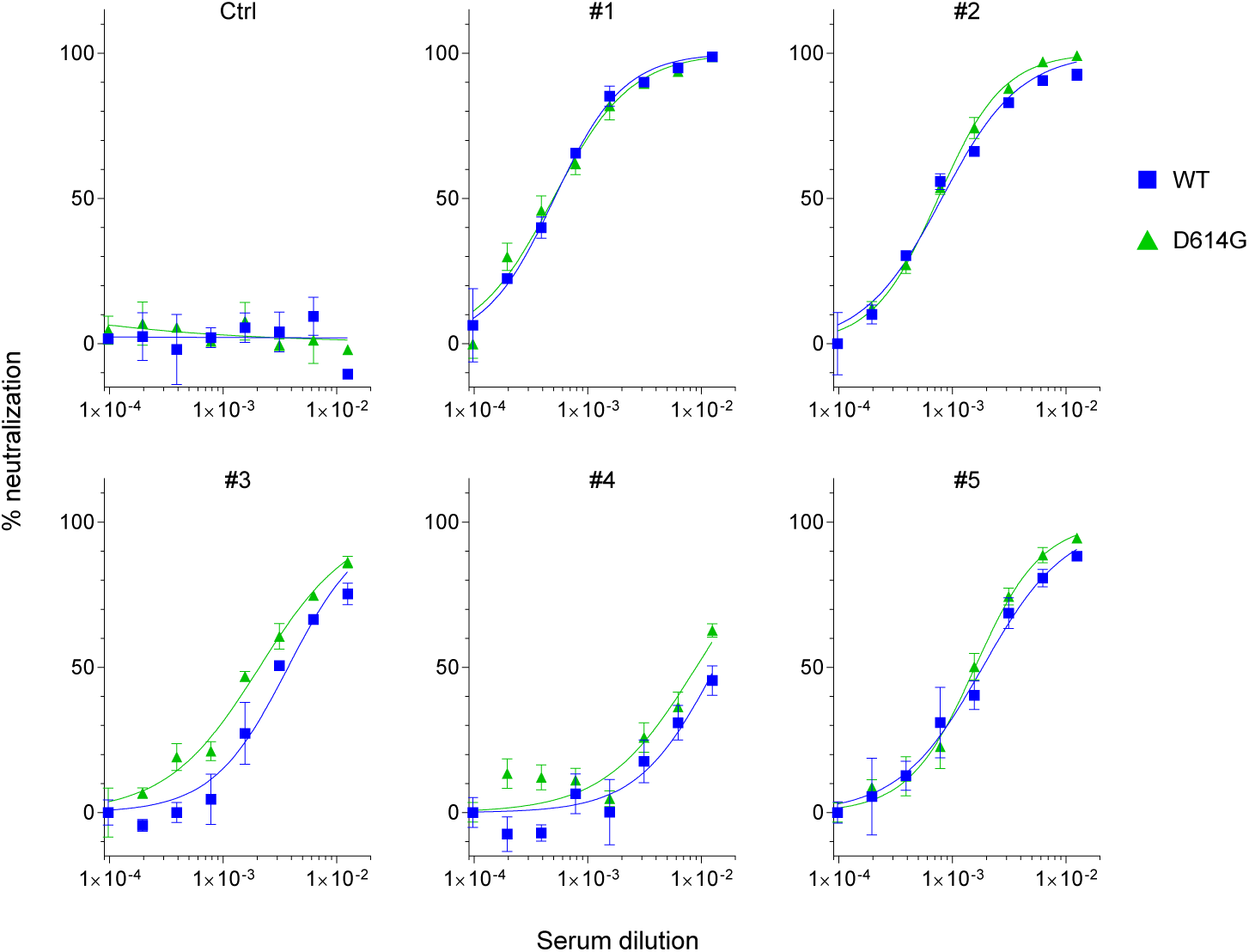
The prototype (wild-type, WT) and D614G mutant SARS2-S are similarly neutralized by patient sera. Lentiviruses pseudotyped with either the WT or D614G mutant SARS2-S were preincubated with two-fold serially diluted human sera (80-fold to 10, 240-fold) obtained from a healthy donor (Ctrl) or collected at 15-30 days post-symptom onset from confirmed case patients (#1 - #5) infected with the prototypic viruses. The mixture was used for infection of 293T cells coexpressing ACE2 and TMPRSS2, and cell entry levels of pseudoviruses in the presence of diluted patient sera were determined by luciferase assays. Representative data from two independent experiments are shown as percent neutralization (mean ± s.d., *n* = 3 technical replicates).

In this study, we employed a novel entry assay system that we recently developed to quantify cell entry of lentiviral pseudoviruses (*10*), in which we can precisely normalize viral input unlike for retrovirus- or vesicular stomatitis virus-based pseudoviruses, leading to experimental accuracy in determining levels of viral entry. By using this system, we first showed that SARS2-S-mediated cell entry strongly depends on TMPRSS2 coexpression, without which that of SARS-S still proceeds to some extent. Indeed, SARS-CoV-2 efficiently infects respiratory and intestinal cells (*9,22*) that coexpress ACE2 and TMPRSS2 (*8*), whereas SARS-CoV can target multiple cell types in several organs (*23*) expressing ACE2, probably even without TMPRSS2. We also found that SARS2-S-mediated cell entry is considerably lower than that of SARS-S, consistent with the recent report that SARS-CoV replicates ∼1,000-fold more efficiently than does SARS-CoV-2 in highly permissive human intestinal Caco2 cells (*24*). These findings suggest the possibility that SARS2-S is likely less adapted to human ACE2 than SARS-S, and might, at least in part, explain why asymptomatic infection is very high in the case of COVID-19, e.g., in a nursing facility in Washington, U.S.A. (56%; 27 out of 48) (*25*), and among healthcare workers in the UK (57%; 17 out of 30) (*26*), whereas it was very rare in the SARS outbreak in Hong Kong in 2003 (0.19%; 2 out of 1,068) (*27*).

Second and more importantly, the D614G mutant displayed the highest entry activity among SARS2-S proteins tested in this study (∼3.5-fold higher than that of the WT protein). Because a single round virus infection is amplified during multiple virus replication cycles, such a seemingly small increase in entry activity can lead to a large difference in viral infectivity *in vivo*. This finding therefore suggests that the D614G mutant virus might be more transmissible from human to human than the prototype viruses, even though further investigations are necessary to determine the correlation between viral infectivity and transmissibility among humans. In fact, the D614G mutation defines the clade A2a, which has overwhelmingly spread worldwide, accounting for 87% of sequenced cases in New York City (73 out of 84 in one study (*20*)), 76% of cases in Iceland (438 out of 577, including A2a-derived haplotypes in a study (*17*)), the vast majority of the second-wave cases in Japan (*19*), and almost all of the cases in Italy (*19*). It should be noted that in 15 European countries where the A2a clade has been dominant, the estimated doubling time was reported to be ∼3 days or less (*28*), which is significantly shorter than the initial estimation (6.4–7.4 days) in the early outbreak phase in China (*2,29*).

Third, the aforementioned results indicate that the D614G variant has biological and structural differences from the WT strain. These findings prompted us to assess whether this mutation might have changed the S protein’s antigenicity, as generally represented by reduced antibody-neutralization susceptibility. According to the results of pseudovirus-based neutralization assays, we found that the SARS2-S WT and D614G mutant were similarly susceptible to all confirmed case patient sera raised against the prototypic viruses. This is particularly important in that the clade-defining D614G mutation will not hinder the current strategy for anti-SARS-CoV-2 drug/vaccine development due to the absence of antigenic alteration in the S protein. It should be noted, however, that all of these infection experiments including viral entry assays were performed by using lentiviral vector-based S-protein-pseudotyped viruses, and therefore the results obtained need to be reproduced by conducting infection experiments using whole SARS-CoV-2 viruses in future studies.

Overall, we speculate that the reason why COVID-19 has become a once-in-a-century pandemic while SARS ended up with epidemics in the past could be, at least in part, that the S protein, a key viral protein for the first step in transmission, of the causative agent SARS-CoV-2, displays suboptimal levels of cell entry activity as shown in this study. Accordingly, this phenomenon would occasionally result in inefficient replication among humans, presumably leading to a higher rate of asymptomatic infection, as described above. In such cases, unknowingly infected subclinical individuals could be virus spreaders who still have the potential to cause lethal respiratory disease to others, as recently reported (*25*). Consequently, during its continuous transmission from human to human, the Wuhan prototype might have acquired more widely prevalent phenotypes represented by the A2a clade that harbors the D614G mutation, possibly with enhanced speed of global transmission. If this scenario is correct, fatal pathogens for which we should be concerned are those with such characteristics rather than the obviously deadly viruses that can be readily detected. Although in the present case, it is likely that the mutation did not influence the antigenicity of the S protein, we may need further worldwide surveillance for virological changes of SARS-CoV-2 in the human population.

## Acknowledgments

We thank the following physicians who collected blood samples from confirmed case patients with COVID-19: Sakiko Tabata, Mayu Ikeda, and Kaku Tamura (Self-Defense Forces Central Hospital). This work was supported by grants from the Japan Society for the Promotion of Science (KAKENHI, 18K07156) to K.T., Kumamoto University Internal Grant for COVID-19 Research to U.T., the intramural research grant of Nagoya Medical Center to Y.I., and the Japan Agency for Medical Research and Development (AMED, Research Program on COVID-19, JP19fk0108110) to T.S.

## Author Contributions

S.O., Y.Z., H.O., T.S.T., T.U., Y.I., and K.T performed experiments and analyzed the data. H.O., S.K., T.U., Y.I., T.S. and K.T. interpreted and discussed the data. K.I., K.M. provided reagents. K.T. conceived the study, supervised the work and wrote the paper. All authors read and approved the final manuscript.

The findings and conclusions in this paper are those of the authors and do not necessarily represent the official positions of the National Institute of Infectious Diseases, University of Yamanashi, National Hospital Organization Nagoya Medical Center, Kumamoto University, and Self-Defense Forces Central Hospital.

## Competing Interests

The authors declare no competing financial interests.

## Data and Materials Availability

Plasmids are available from K.T. under a material transfer agreement with the National Institute of Infectious Diseases.

## Materials and Methods

### DNA constructs

The HiBiT-tagged lentiviral packaging plasmid psPAX2-IN/HiBiT, the firefly luciferase-expressing lentiviral transfer plasmid pWPI-Luc2, CMV/R-SARS-S, and the ACE2-expressing plasmid have previously been described elsewhere (*11, 30, 31*). The ACE2 expression plasmid pC-ACE2 was created by inserting the Acc65I/XhoI-digested PCR-amplified human ACE2 fragment into the corresponding site of the pCAGGS mammalian expression plasmid (*32*). To generate the TMPRSS2 expression plasmid pC-TMPRSS2, total RNA isolated from HepG2 cells was subjected to RT-PCR amplification of the TMPRSS2 gene using specific oligonucleotides, and an amplified fragment digested with BsiWI/XhoI was cloned into Acc65I/XhoI-digested pCAGGS (possible RT-PCR errors or SNPs were corrected by PCR mutagenesis). The SARS-CoV S expression plasmid pC-SARS-S was created by inserting the BsiWI/XhoI-digested PCR-amplified SARS-CoV S fragment of CMV/R-SARS-S into the corresponding site of pCAGGS. The SARS-CoV-2 S expression plasmid pC-SARS2-S was created by inserting the Acc65I/NotI-digested PCR-amplified SARS-CoV-2 S fragment of pCMV3-2019-nCoV-Spike(S1+S2)-long (Sino Biological; VG40589-UT) into the corresponding site of pCAGGS. The SARS-CoV-2 S mutants (pC-SARS2-S-H49Y, pC-SARS2-S-V367F, pC-SARS2-S-G476S, pC-SARS2-S-V483A, pC-SARS2-S-D614G, or pC-SARS2-S-C1247A), in which positions 49, 367, 476, 483, 614, or 1247 of the S protein were mutated from histidine to tyrosine, valine to phenylalanine, glycine to serine, valine to alanine, aspartic acid to glycine, or cysteine to alanine, respectively, were created by inserting overlapping PCR fragments into Acc65I/NotI-digested pCAGGS. The plasmid expressing chimeric SARS2-S protein harboring the TM/CT domains of SARS-S, pC-SARS2-S-TM/CT1, was generated by inserting overlapping PCR fragments (amplified from the TM/CT domains of SARS1-S and from all other domains of SARS2-S) into Acc65I/NotI-digested pCAGGS. All constructs were verified by a DNA sequencing service (FASMAC).

### Cell maintenance and stable cell establishment

293T and HepG2 cells were maintained under standard conditions. Cells were originally obtained from the ATCC and routinely tested negative for mycoplasma contamination (PCR Mycoplasma Detection Set, Takara).

### Pseudotyped virus preparation and cell entry assays

To prepare various spike protein-pseudotyped lentiviral luciferase reporter viruses, 1.1 × 10^5^ 293T cells were cotransfected with 200 ng of spike protein expression plasmids (pC-SARS-S, pC-SARS2-S (-WT, -H49Y, -V367F, -G476S, -V483A, -D614G, or -C1247A), 400 ng of psPAX2-IN/HiBiT, and 400 ng of pWPI-Luc2, using FuGENE 6 (Promega). Sixteen hours later, the cells were washed with phosphate-buffered saline, and 1 ml of fresh complete medium was added. After 24 h, the supernatants were harvested and treated with 37.5 U/ml DNase I (Roche) at 37°C for 30 minutes. The lentivirus levels in viral supernatants were measured by the HiBiT assay, as previously described (*10*). Briefly, a lentivirus stock with known levels of p24 antigen was serially diluted. Either the standards or viral supernatants containing pseudotyped viruses (25 μl) and LgBiT Protein (1:100)/HiBiT Lytic Substrate (1:50) in Nano-Glo HiBiT Lytic Buffer (25 μl) (Nano-Glo HiBiT Lytic Detection System; Promega) were mixed and incubated for ten minutes at room temperature according to the modified manufacturer’s instructions. HiBiT-based luciferase activity in viral supernatants was determined with a Centro LB960 luminometer (Berthold). To prepare target cells, 2.2 × 10^5^ 293T cells were cotransfected with increasing amounts of pC-ACE2 and pC-TMPRSS2, together with 700 ng of empty pCAGGS plasmid. After 48 h, transfected cells (2.2 × 10^4^) were seeded into 96-well plates, and incubated with lentiviruses pseudotyped with different spike proteins corresponding to 1 ng of p24 antigen. After 48 h, the cells were lysed in 100 μl of One-Glo Luciferase Assay Reagent (Promega). Firefly luciferase activity was determined with a Centro LB960 luminometer. Note that in this procedure, HiBiT assays using target cell lysates were not employed to determine the levels of cell entry, due to the background caused by non-specific attachment of virions to the cell surface.

### Spike incorporation assays

Supernatants containing either SARS2-S-pseudotyped viruses were layered onto 20% (wt/vol) sucrose cushions and subjected to ultracentrifugation (90,000 r.p.m. for 10 minutes) using an Optima TLX Ultracentrifuge (Beckman Coulter). Pelleted virions were resuspended in SDS-sample buffer and subjected to immunoblot analysis using an anti-SARS-CoV-2 spike mouse monoclonal antibody (1:1,000, GeneTex, GTX632604) and an anti-p24 monoclonal antibody (1:1,000; Nu24 (*33*)).

### Structural modeling of SARS-CoV-2 S proteins

The structure-based complex models of ACE2 and the ectodomains (ECs) of SARS-CoV-2 S proteins were built by homology modeling with the Modeller 9v8 (*34*). The trimeric structure of the EC (PDB code: 6VYB (*35*)) and that of the RBD-ACE2 complex (PDB: 6M0J (*36*)) obtained from the Protein Data Bank, and was used as a template for the modeling. The GenBank reference sequences of SARS2-S and human ACE2 were used for the modeling (GenBank IDs: YP_009724390.1 and NP_001358344.1). The structural models of the H49Y, V367F, G476S, V483A, and D614G mutants were constructed in a similar manner. The complex model of ACE2 and SARS-CoV S’s EC was also constructed by using the structures of the EC trimer (PDB: 6CS1 (*37*)), its RBD binding to ACE2 (PDB: 3D0G (*38*)), and the reference SARS-S sequence (NP_828851.1). The structural figures were generated by using PyMOL ver. 2.4 (https://pymol.org).

### Neutralization assays

Experiments using human samples were approved by the Medical Research Ethics Committee of the National Institute of Infectious Diseases, Japan. Five serum samples were collected at 15-30 days post-symptom onset from confirmed case patients (obtained in February 2020) and heat-inactivated at 56°C for 30 minutes. Two-fold serially diluted sera were mixed with an equal volume of 1 ng of 24 antigen of the WT or D614G mutant SARS2-S-pseudotyped virus and incubated at 37°C for 1 h. The mixture was added to 293T cells transiently coexpressing ACE2 and TMPRSS2 (seeded into a 96-well plate). After 48 h, cells were lysed and subjected to luciferase assays, as described above, to determine the levels of neutralization.

### Statistical analyses

Values are presented as the mean ± s.d. for three to seven independent experiments determined on the basis of pilot experiments to estimate effective numbers. Statistical analyses of the data were performed by using GraphPad Prism version 8.04. Statistical comparisons were made using a two-tailed paired Student’s *t*-test or one-way ANOVA with Dunnett’s multiple comparison test.

---

**Figure S1.**
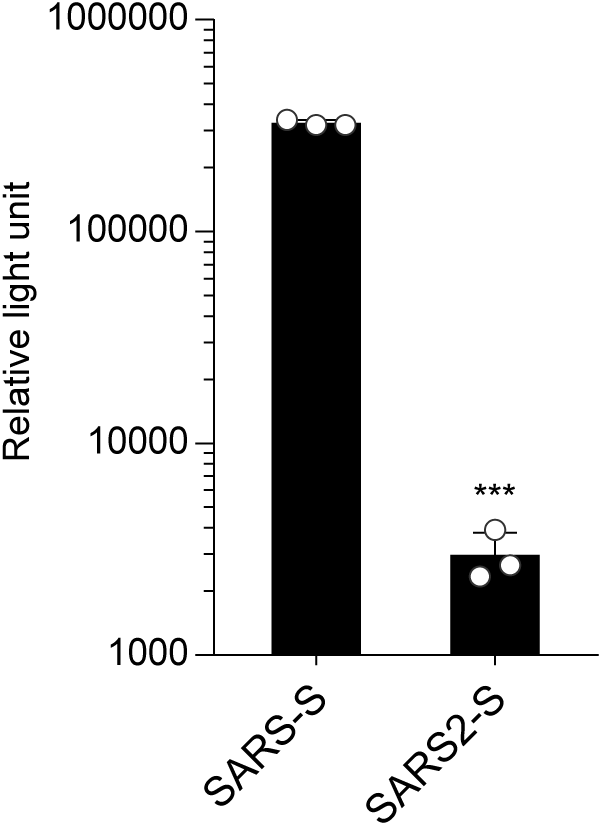
ACE2 expression alone is insufficient to support SARS2-S-mediated cell entry. To prepare S-pseudotyped lentiviruses, 293T cells were transfected with the HiBiT-tagged lentiviral packaging plasmid, the firefly luciferase-reporter lentiviral transfer plasmid, and either a SARS-CoV S (SARS-S) or SARS-CoV-2 S (SARS2-S) expression plasmid. The viruses produced were assessed by HiBiT assays, and S-pseudotyped viruses normalized based on HiBiT activity were used for infection of 293T cells expressing the host receptor ACE2 only. Cell entry was determined by firefly luciferase activity in cell lysates. Data from three experiments are shown (mean ± s.d., *n* = 3 technical replicates). The *p* value was calculated using paired two-tailed Student’s t-test, \*\*\**p* < 0.001.

**Figure S2.**
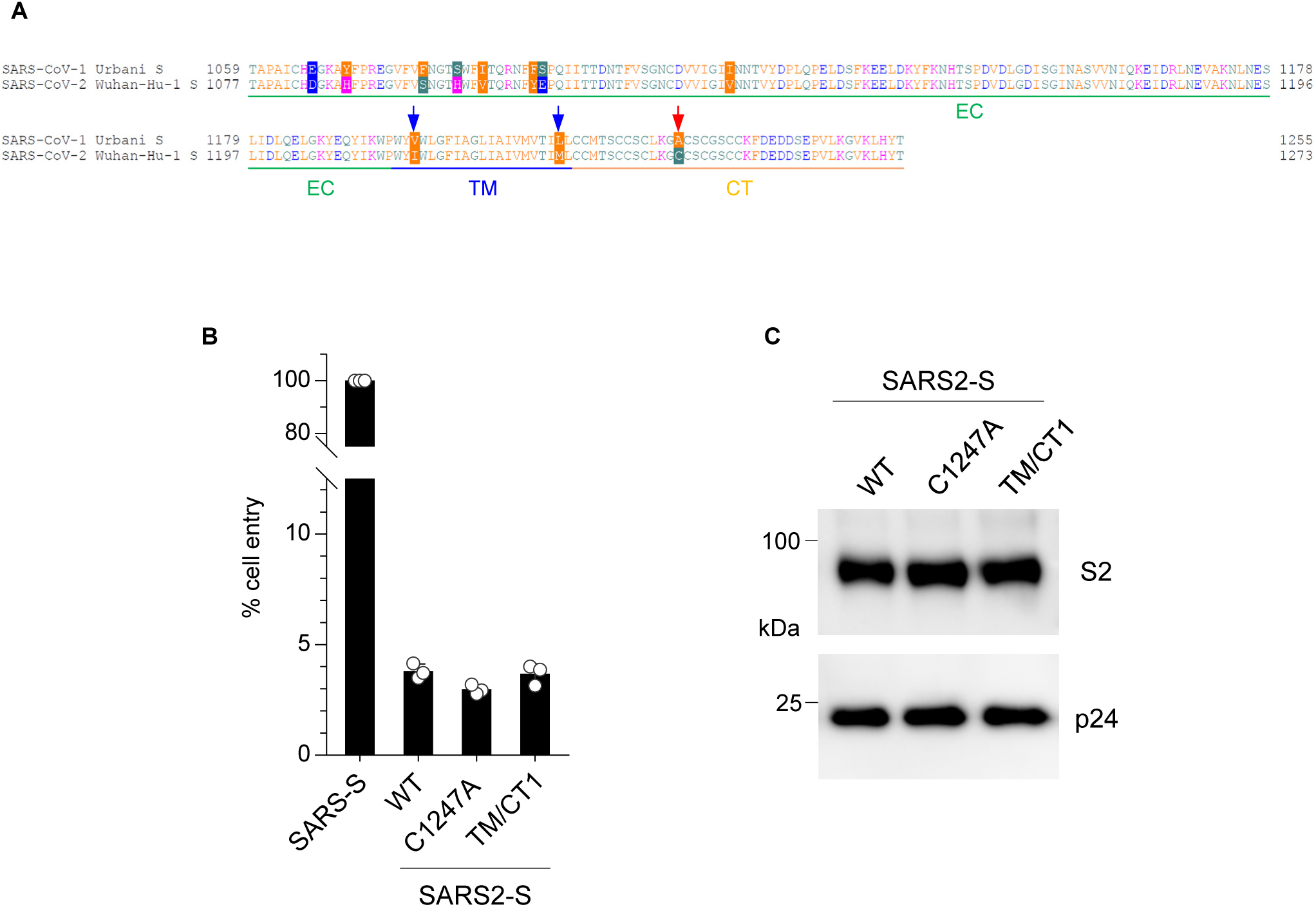
The lower level of SARS2-S-mediated entry is not due to differences in the efficiency of S incorporation into virions. (A) Amino acid sequence alignments of the C-terminal sequence of S proteins from SARS-CoV (Urbani strain) and SARS-CoV-2 (Wuhan-Hu-1 strain). Amino acid differences are boxed in several colors. EC, extracellular domain; TM, transmembrane domain; CT, cytoplasmic tail. SARS2-S-C1247A and SARS2-S-TM/CT1 were created by mutating a cysteine to an alanine at the CT (indicated by a red arrow) and by additionally mutating an isoleucine and a methionine to a valine and a leucine at the TM (indicated by blue arrows), respectively. (B) 293T cells were transfected with the HiBiT-tagged lentiviral packaging plasmid, the firefly luciferase-reporter lentiviral transfer plasmid, and the plasmid expressing either the SARS2-S wild-type (WT) protein or one of the two mutants (SARS2-S-C1247A or SARS2-S-TM/CT1). Viruses produced were assessed by HiBiT assays, and S-pseudotyped viruses normalized based on HiBiT activity were used for infection of 293T cells expressing ACE2 and TMPRSS2. Cell entry was determined by firefly luciferase activity in cell lysates. Data from three experiments are shown as a percentage of cell entry of the SARS-S-pseudotyped viruses (mean ± s.d., *n* = 3 technical replicates).

## Notes

### Competing Interest Statement

The authors have declared no competing interest.

### Summary of Updates

An author's name was corrected, the references were amended, and Abstract and Methods were slightly revised.

